# Age-Related Differences in the Lexical Priming Effect Can Be Predominantly Attributed to Differences in Response Bias

**DOI:** 10.64898/2025.12.04.689606

**Authors:** Mo Eric Cui, Bruce Schneider, Cristina D. Rabaglia

## Abstract

Age-related declines in visual perception challenge older adults’ ability to locate and identify objects in noisy (e.g., fog, glare, inadequate lighting) or cluttered visual environments. To compensate, older adults often increase reliance on contextual information, which could be helpful, but may also lead to increased risk for “false seeing” – reporting expected objects based on contextual information rather than those actually presented – a phenomenon that Jacoby et al., (2012) observed in older adults under challenging viewing conditions. However, it remains unaddressed whether such false seeing is driven by older adults’ elevated tendency to rely on context, or by age-related declines in visual perceptual processes. The current study addressed this issue by examining the lexical priming effects in 18 younger and 18 older adults (*M* = 21.4 years; *M* = 76.2 years) in Experiment 1, and 13 younger and 13 older adults (*M* = 21.2 years; *M* = 74.8 years) in Experiment 2 using a two-alternative, forced-choice (2AFC) paradigm, while controlling for age-related perceptual differences. On each trial, participants identified a visually degraded target word preceded by an identical, semantically related, or unrelated prime. We applied Signal Detection Theory to isolate perceptual sensitivity (*d*’) and response bias (*c*). Results revealed that once stimulus parameters were adjusted to eliminate age-differences in performance when the prime was unrelated to the target, age-related differences in target identification in the related priming conditions were predominantly driven by age-related differences in response bias rather than in any residual age-differences in perceptual sensitivity.

**Public Significance Statement:** To compensate for age-related losses in visual processes, older adults tend to rely more on the overall context provided by the visual scene to identify individual objects than do younger adults. Using a word priming task, our study found that older adults’ use of supporting context to correctly identify target words masked by visual noise persisted even when overall word identification performance (correctly identifying words independent of whether the context was supporting or misleading) was equated for both age groups. Understanding such age-relate differences in visual perception can inform interventions that would support older adults’ visual processing in daily perceptually challenging environments.

---

Age-related visual declines make it difficult for older adults to identify objects under difficult viewing conditions (e.g., inadequate lighting, fog, visual clutter) forcing them to rely more heavily on contextual information to organize the visual field and identify the objects in it. Under such conditions, compared to younger adults, older adults are more often report *seeing* expected, but incorrect, objects – a phenomenon termed “false seeing” (Jacoby et al., 2012). In this study, we use Signal-Detection Theory (SDT) to determine the degree to which age-related differences in perceptual acuity and/or in response bias, can account for age-related differences in “false seeing.”

Aging is associated with declines in visual perception, including reduced contrast sensitivity and visual acuity (Bennett et al., 2007; Owsley, 2011; Owsley et al., 1983; Sekuler & Sekuler, 2024; World Health Organization, 2019). These changes arise from both ocular factors - such as pupil constriction, lens hardening, and cataract formation (Owsley, 2011; Owsley et al., 1983; Weale, 1983) and neural degeneration in the retina and primary visual pathways (Sekuler & Sekuler, 2024). As a result, older adults face greater perceptual challenges than younger adults especially when they encounter visually demanding environments in daily life (e.g., a foggy day). In such situations, on textual information can serve an important compensatory role, allowing older adults to mitigate the effects of perceptual deficits through top-down processing in many scenarios (Pichora-Fuller et al., 1995; Schneider & Pichora-Fuller, 2000; Speranza et al., 2000). For example, when viewing a blurry or partially obscured image, a mountain forest background might facilitate recognizing a somewhat obscured image as a gorilla, when compared to a presentation of same obscured image in a cityscape.

## Context as a Compensatory Strategy

Numerous studies have demonstrated that older adults use contextual information to support perception (Faulkner, 1983; Pichora-Fuller et al., 1995; Schneider & Pichora-Fuller, 2000). Auditory recognition studies consistently show enhanced contextual benefits among older listeners using manipulations like time-compressed speech (Moberly et al., 2023; Sheldon et al., 2008), multi-talker babble (Goy et al., 2013; Rogers et al., 2015), and low-pass filtering (Goy et al., 2013). An increased reliance on contextual information in older adults has also been noted in visual recognition studies (Madden, 1988; Rayner et al., 2009; Speranza et al., 2000). For example, in lexical priming tasks, the observer is asked to identify a target word in a visually noisy background. In such tasks the target word is preceded by a clearly visible ‘priming word’ that may or may not offer relevant contextual cues to facilitate the recognition of the target word. Generally, related primes improve recognition accuracy compared to unrelated primes, and the literatures indicate that older adults rely more heavily on these contextual primes than younger adults (Laver & Burke, 1993). Notably, older adults often show the *hyperpriming* effect, an exaggerated facilitation effect for strongly associated pairs of words, which is likely related to their increased reliance on preserved semantic memory networks (Bowles & Poon, 1988).

Although the literatures listed above supports greater contextual reliance in older adults, not all studies have found such age-related differences in the effects of context. For example, some studies reported comparable semantic benefits between younger and older adults during a speech recognition task (Dubno et al., 2000), while other studies suggested that older adults may be less effective at forming predictive expectations from contextual cues (Federmeier et al., 2005). Dubno et al. (2000) argued that such inconsistent findings might be due to differences in baseline perceptual difficulty that can either enhance or suppress the measured contextual effects. Studies equating perceptual difficulties at a group level fail to consider individual variation in perceptual processes, which tend to increase with age, and may not be as responsive to age-related differences in visual perception as are those studies that adjust for perceptual difficulty at an individual level.

## Misleading Context and False Seeing

Beyond supporting recognition, contextual cues may sometimes mislead perception. If a blurred image is presented in a misleading scene (e.g., an obscured gorilla in a cityscape), viewers may mistakenly identify the target as something congruent with the context, like a person. Given their visual deficits and reliance on context, older adults may be especially vulnerable to this error, a phenomenon named “false seeing” (Jacoby et al., 2012). Indeed, Jacoby et al., (2012) showed that older adults were more likely to mistakenly report seeing a misleading prime (e.g., “DART”) rather than the actual, briefly flashed target (e.g., “dirt”). Similar effect has been observed in auditory research under misleading sentential contexts, suggesting potential deficits in inhibitory control or heightened susceptibility to interference (Failes & Sommers, 2022; Rogers & Wingfield, 2015).

A critical question is the extent to which this “false seeing” phenomenon reflects can be attributed to age-related perceptual declines. Jacoby et al., (2012) attempted to control for baseline perceptual difficulty by individually adjusting target presentation duration to account for age-related slowing of processing speed (Salthouse, 1996). However, this method failed to directly control for persistent visual deficits, such as reduced contrast sensitivity and visual acuity, which are independent of stimuli duration. Consequently, older adults heightened reliance on context may be a compensatory response to unaddressed sensory impairments, rather than a genuine cognitive bias.

A potentially more effective approach would screen participants for poor vision and adjust stimulus contrast (signal-to-noise ratio, SNR) separately for younger and older adults to achieve matched baseline accuracy (e.g., 75% correct identification in 2AFC) in the absence of contextual cues. Equating accuracy across age groups in the absence of contextual cues thereby provides a baseline reference against which to evaluate age-related differences when contextual cues are introduced. Individuals may also differ in the extent to which they use context to identify objects in their daily lives. For example, visually-impaired individuals may rely more on context to parse difficult-to-see objects, much as observers of the blurry “gorilla” image used the urban scene context to bias their interpretation toward “human.”

## A Signal Detection Theory (SDT) Approach

To clarify whether age-related differences in context reliance stem from cognitive bias or perceptual sensitivity, we employed a novel application of SDT. Typically, SDT differentiates perceptual sensitivity (*d’*) from response bias (*c*). The sensitivity of the observer to the presence of the signal in the noise is assessed by *d’*. Response bias represents the criterion individuals adopt to make decisions under uncertainty. While SDT paradigms often involve a detection task (signal-plus-noise vs. noise-alone), the current experiment uses a 2AFC word identification task. Experiment 1 established individualized stimulus contrast levels to equate perceptual sensitivity (*d′*) across age groups in a word identification task where the prime word was unrelated to the target word. Experiment 2 then utilized a novel SDT approach to determine if age-related differences in performance were driven by changes in response bias rather than age-related sensory deficits.

The SDT analysis employed here distinguishes between two types of trials. In both trial types a lower-case target word is presented in visual noise. In a Prime = Target type of trial, prior to the presentation of the lower-case target word in noise, a clearly visible priming word is presented in upper-case letters against a uniform gray background (e.g., DIRT/dirt). In a Prime = Foil trial, a clearly visible priming word that is related to, but differs from the target word by one letter, is presented in upper-case letters before the lower-case target word is presented in noise (e.g. DART/dirt). In both trial types, after the presentation of the target in noise, the participant is visually informed that the target word that was just presented was either “dirt” or “dart”. Note that in the SDT analysis conducted here, there is no noise-alone trial type. Hence, in the SDT analysis of this paradigm, we define the decision axis as the amount of evidence supporting a decision that the target is equal to the prime (Figure 1).

**Figure 1.**
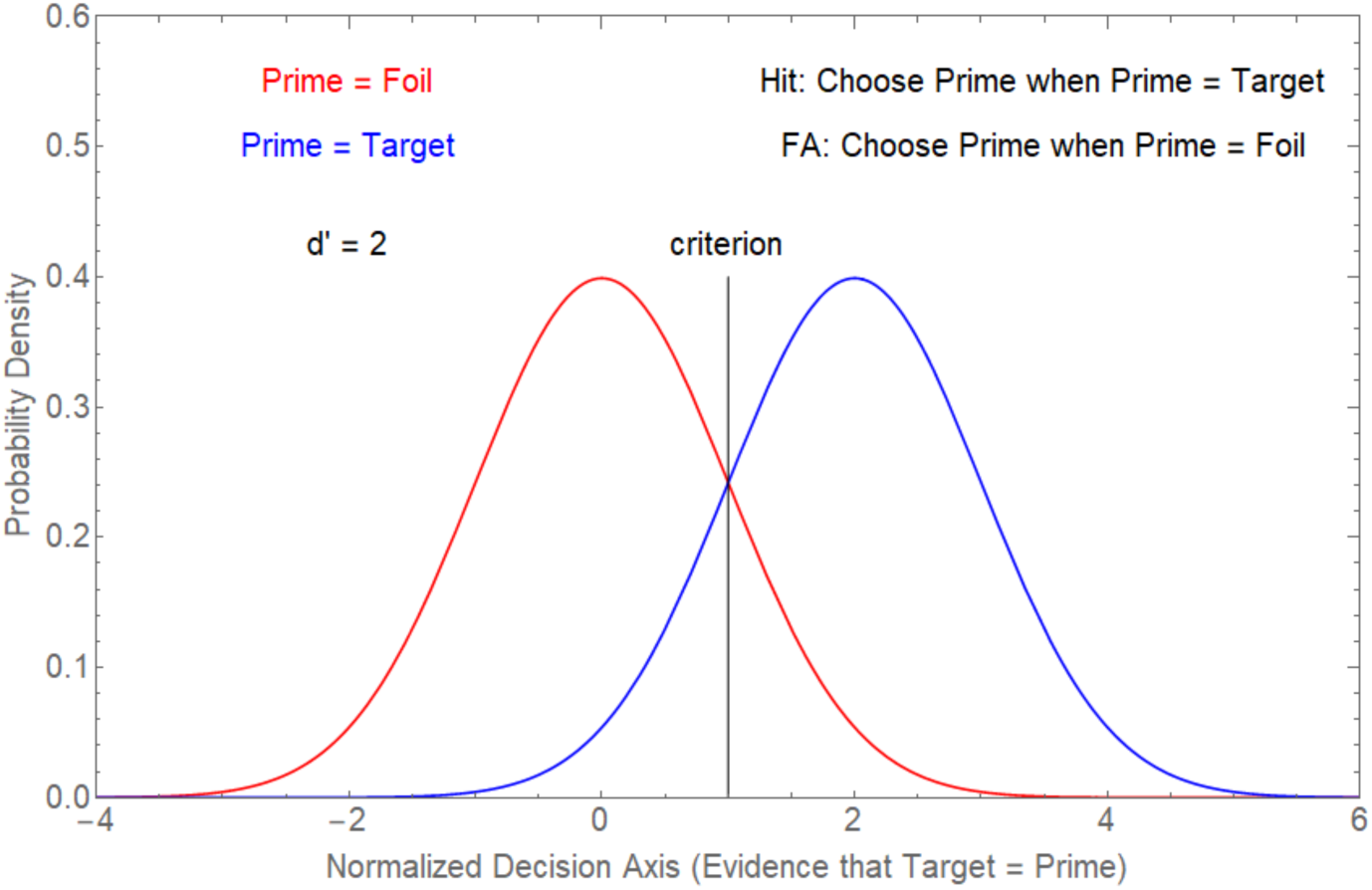
Signal Detection Model of a Priming Experiment. Note. The decision axis represents the evidence that the target is the Prime. The ideal observer would gather evidence that the Target was the same as the Prime. The blue curve represents the normalized distribution of the amount of such evidence when prime = target. The red distribution represents the normalized distribution of the amount of evidence that the target is the prime when the prime = foil. The discriminability between the two distributions (*d’*) is the normalized distance between the peaks of the two distributions. Because the target was presented in noise, the amount of evidence gathered during a noisy target presentation would vary from trial to trial, with the amount of evidence being more for Target = Prime trials than for Target ≠ Prime trials. If the amount of evidence exceeds the criterion amount selected by the observer, the observer identifies the target as being identical to the prime. When the probability of both trial types is 0.5, correct identifications of target are maximized when the criterion is located at the intersection of the two distributions. In this Figure, *d’* was set to 2.

In this interpretation of a priming experiment, the observer, on a trial, accumulates evidence that the target is the same as the prime. More evidence that the target is the same as the prime accumulates when the target is the same as the prime, than when the target is not equal to the prime. Due to visual noise, the amount of evidence accumulated is noisy, and is assumed to vary in a Gaussian fashion from trial to trial. The observer establishes a criterion along this decision axis: if this criterion is exceeded, the observer identifies the target as the prime, otherwise the target is identified as the foil. Thus, we can directly examine whether younger and older adults differ in how they weight and utilize contextual evidence during target identification within a signal-detection framework. Hence, if older adults depended on context more than younger adults, result would be that their decision criterion would be located to the left of the unbiased position shown in Figure 1, while the location of the criterion for younger adults would be to the right of the location of the criterion used older adults.

## The Current Study

To recapitulate, the current study was designed to address the following question: when the stimulus situation in a priming experiment is adjusted to produce equivalent discriminability, do older adults rely more on contextual cues provided by the prime than do younger adults? We looked for age-related differences in priming effects, after adjusting the stimulus situation to produce equivalent discriminability in both age groups and used the signal-detection model described in Figure 1 to assess the extent to which the previously observed ‘false seeing’ was driven by age-related difference in the use of contextual information, or actual deficits in sensory processing. We used the results from Experiment 1, where the prime was unrelated to the target, to determine signal-to-noise ratios (SNRs) that would result in equivalent performance in younger and older adults in the absence of any valid information in the prime that could improve performance when the target was presented in visual noise. Then, in Experiment 2, where the prime was either the same as the target (Prime = Target), or related but different from the target (Prime = Foil), we used a SDT (Figure 1) to assess the contribution of age-related differences in response bias to age-related differences in ‘false seeing.

## Method

### Participants

Older participants were recruited from the University of Toronto Mississauga Senior Volunteer Pool. Younger participants were recruited from the university’s undergraduate population through the SONA system. Eligible participants self-identified as native English speakers and reported no history of severe psychological, neurological, or vision-related disorders. Participants with a history of traumatic brain injury or recent vision surgery (i.e., in the last 6 months) were excluded. The study received ethical approval from the University of Toronto’s Office of Research Ethics (ORE). Participants received a small remuneration for their participation.

### Measures

#### Visual Assessment

Visual acuity was assessed using the LogMAR scale, and contrast sensitivity was measured with the F.A.C.T. (Functional Acuity Contrast Test) across various spatial frequencies using an Optec 6500P device (Stereo Optical Company, Chicago, IL). Participants wore the appropriate corrective lenses during this visual acuity assessment and during the experiment itself. These measures assessed and confirmed typical age-related visual differences between groups and characterized participants’ sensory profiles.

#### Mill Hill Vocabulary Test

Participants completed the Mill Hill Vocabulary Test (Raven, 1965), a 20-item forced-choice test assessing English proficiency and vocabulary size. Test items were presented with six alternatives, and participants had to select the closest synonym to a test item. There was no time constraint. The total score was calculated as the number of correct responses out of 20. This measure was included to characterize participants’ general verbal knowledge, and this measure has been commonly used as an index of cognitive abilities.

### Materials

The experimental materials included 180 triplets of common four- and five-letter English words, and were selected based on the criteria described in Jacoby et al. (2012). Each triplet consisted of two similar words (i.e., a similar word pair) differing by a single critical letter at the same position (e.g., *dirt* – *dart*, *critical location* = 2) and an unrelated word. The unrelated word neither contained the critical letter nor shared any letter in the same position with the similar pair (e.g., *dirt* & *dart* – *chew*; or *cold* & *bold* – *dark*). Four-letter word pairs could differ at any one of 4 critical locations, and five-letter word pairs could differ at any one of 5 critical locations. Thus, we carefully selected 20 similar word pairs for each one of these 9 critical location groups for a total of 180 similar word pairs. Then, we assigned an unrelated word for each pair, making a total of 180 word-triplets. We controlled some of the key lexical characters across these 9 groups. Word frequency ranged from 2.6 to 2.82 counts per 50 million TV subtitles (SUBTLEXUS database; Brysbaert & New, 2009). Orthographic neighbor counts, defined as the number of words differing by only one letter (MRC Psycholinguistic Database), only differed between the four-letter groups and five-letter groups (*M =* 13.11, *M =* 6.93; *t*(178) = 12.89, *p* < .001). No significant differences were found within the 4 four-letter or the 5 five-letter groups (*F*s < 1).

### Experimental Design and Procedure

The experimental design, adapted from Jacoby et al. (2012), consisted of three within-subject conditions: *prime unrelated to target*, *prime = target*, and *prime = foil.* Experiment 1 consisted of 180 baseline trials in which the prime was unrelated to the target. Experiment 2 consisted of 60 trials for each of the three types of prime. At the beginning of the session and between each trial, a 333 × 333 pixel image of a square (11.86° × 11.86°) was projected on the screen in front of the participant. The square had a black border 4 pixels wide (pixel luminance = 255), with all subsequently presented fields having the same black border. The field within the black border was a uniform gray (pixel luminance = 127). Each trial began with a centrally-located black fixation square (4 × 4pixels) displayed for 850 ms, followed by an uppercase prime word presented in black (pixel luminance of the letters = 255, with the letters in the word being 32 pixels tall) for 1 second in the otherwise gray field (pixel luminance = 127). The width of the rectangular area enclosing a letter depended on the letter being displayed, being largest for a capital W and smallest for a capital I. The width of the rectangular area enclosing a word also depended on the number of letters being displayed, ranging from 3.13 to 4.13°, with the height of this area set to be 1°. After a visual mask enclosed by the same black borders (digital Gaussian noise, mean pixel luminance = 127, standard deviation = 40 pixel-luminance levels) lasting 1.5 seconds, a lowercase target word was added to the masker for 850 ms in the center of the masker (mean pixel luminance = 127, root mean square = 40). Specifically, the words in the visual noise were created by adding either 11, 22, 33, or 44 pixel levels to the background visual noise (the higher the pixel level, the darker the luminance of the pixel). Note that the pixels in the letter retained their noisy characteristic. Hence, this had the effect of changing the mean pixel luminance level of the letters in a word to either 116, 105, 94, or 83, without changing the RMS levels of the pixels in the word. We refer to these increments (11, 22, 33, or 44) as decrements in the mean pixel luminance of the noisy letters. Figure 2 presents an example of a word (pout) in which the average pixel luminance of the pixels comprising the letters of the word is 94 (a decrement of 33) but where the RMS level of the pixels in the word is the same as the RMS of the background noise field.

**Figure 2.**
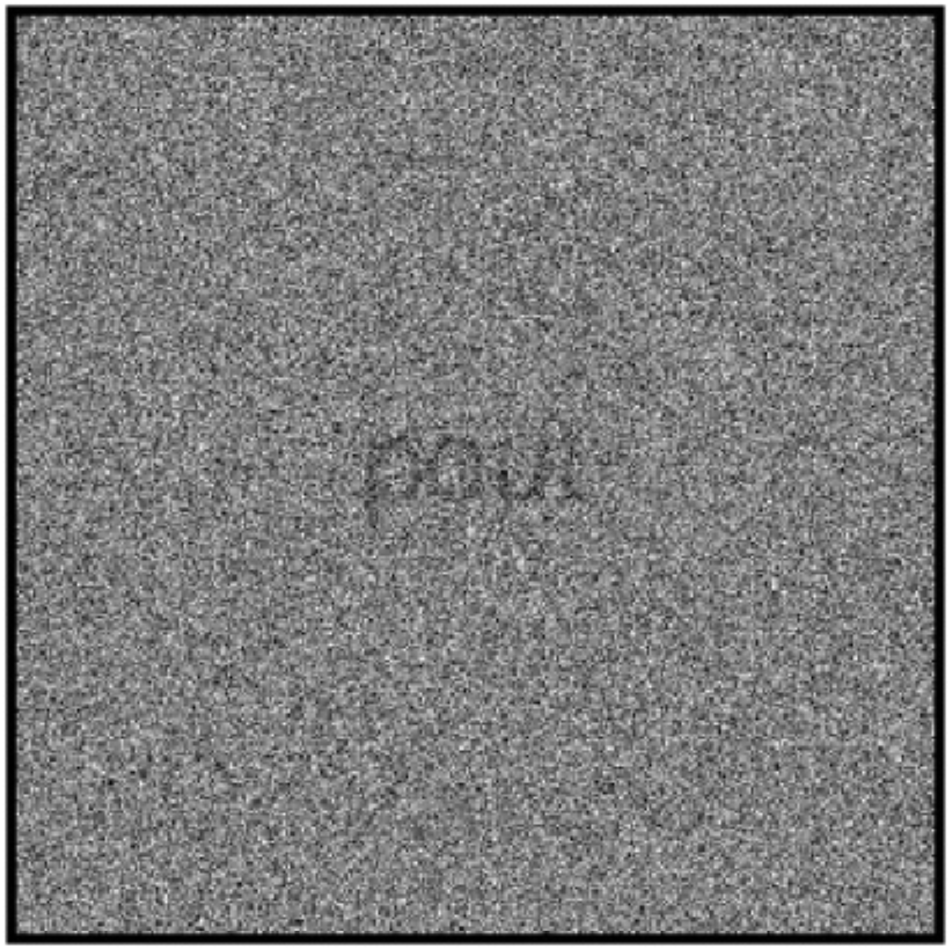
Exemplar Word in Noise Stimuli. *Note. The word ‘pout’ appears in a visual noise background whose mean pixel luminance value is 127 and whose root mean square (RMS) value is 40.* The mean pixel luminance value of the pixels comprising the letters in the word is 33-pixel luminance values lower than that of the background noise) but retains the same RMS value (40) as the pixels in the background noise.

A second independent mask (same characteristics as that of the first mask) followed the lower-case word target for 300 ms, after which participants completed a 2AFC task and a confidence-rating test. In the 2AFC task, participants were informed, after the presentation of the target word in noise, that the target word was one of two possible words that differed only by one letter (e.g., *dart* or *dirt*), presented in random order in lowercase. They then had to choose one of these two words as the target. In the confidence rating task, participants rated their confidence in their performance in this task on a scale from 0 to 5, with 0 representing no confidence, and 5 representing full confidence (See Figure 3).

**Figure 3.**
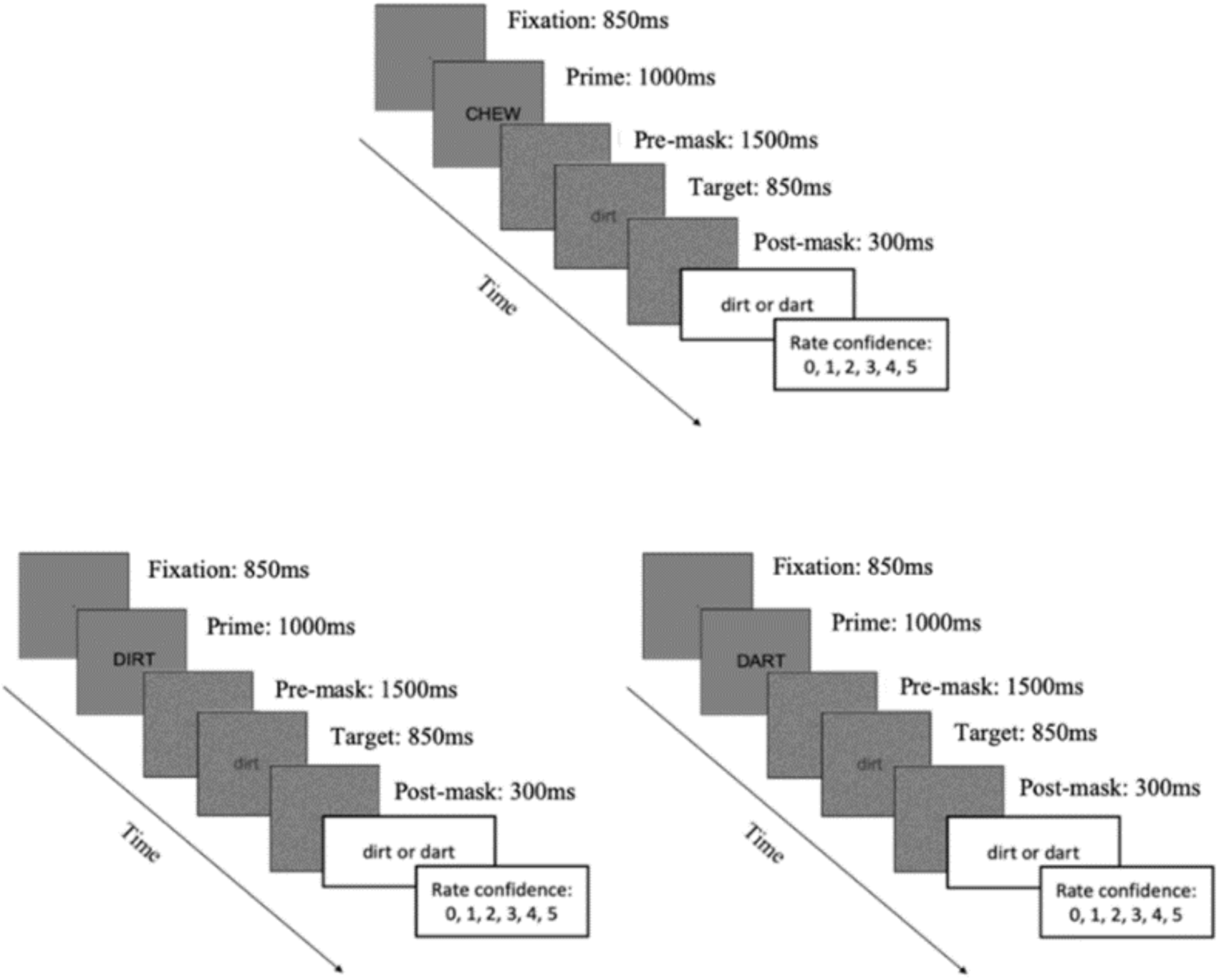
Schematic of Task. *Note*. This figure consists of three lexical priming conditions: unrelated prime (top), prime = target (bottom-left), and prime = foil (bottom-right). Each trial began with a fixation square (850 ms), followed by an upper-case prime word (1000 ms), a pre-mask (1500 ms), a lower-case target word (850 ms), and a post-mask (300 ms). Participants were then informed that the target word was one of two words differing only in a single letter (e.g., “dirt or dart”) and asked to identify which of these two words was the target. They then rated their confidence on a 0–5 scale. In the unrelated prime condition, the prime was a word (e.g, CHEW), that was orthographically and semantically unrelated to the target and foil words (e.g., dirt and dart). In the prime equal target condition, the upper case prime (e.g., DIRT), was semantically identical to the lower-case target (e.g., dirt) and orthographically related to the foil (e.g., dart). In the prime equal foil condition, the upper-case prime (e.g., DART), was semantically identical to the lower-case foil and orthographically related to the target (dart).

In an unrelated prime trial, the priming word was semantically and orthographically different from the target word, and the target word was one of a pair of lower-case words that differed from one another in only one of the four or five letters (e.g., prime = *CHEW*, target = *dirt*, question = *dirt* or *dart*?). In a prime = target trial, the prime word was one of a pair of similar words, one of which was randomly selected as a target (e.g., prime = *DIRT*, target = *dirt*, question = *dirt* or *dart*?). In a prime = foil trial, the prime word was again one of the similar pair of words, and the target was the other word in the pair (the foil). For example, the prime could be *DART*, with the target being *dirt*, and the participant had to choose between either *dirt* or *dart*. In Experiment 1, the extent of the decrement in the mean pixel luminance of the target word was manipulated to determine the function related the probability of correctly identifying the target word to the decrement in mean pixel luminance of the target word when the priming word was unrelated to the target (unrelated prime). The two kinds of related prime (prime = target, prime = foil) were not included in Experiment 1. Experiment 2 used the contrast levels established in Experiment 1 and included all three conditions (unrelated prime, prime = target, and prime = foil).

Participants provided consent and completed a vision screening, demographic questionnaire, and the Mill Hill Vocabulary Test before beginning the lexical priming task. After a brief instruction concerning the word identification task, participants completed a short 10-trial practice block. Participants sat in front of a DELL Inspiron 660 desktop computer and a 16-in. CRT SONY GDM F520 screen (1280 pixel by 1024 pixel) in a dimly lit, double-walled, sound-attenuating booth. Participants’ head position was stabilized with a chinrest placed 18-in away from the computer screen to the center of the screen. The size of the squares presented on the screen was 3.74-in by 3.74-in. The main task consisted of 180 trials divided into three blocks, with an optional mid-session break. The experiment took approximately one hour to complete.

### Transparency and Openness

The experiment materials, code and data are available upon request. Data were preprocessed using custom R scripts (Version 4.4.3) and Wolfram Mathematica-13. This study was not preregistered. Data collection completed by February 2020.

## Experiment 1

### Method

#### Participants

Eighteen younger (*M* = 21.40 years, *SD* = 1.50, range: 19-24 years; 13 females, 5 males) and 18 older adults (*M* = 76.20 years, *SD* = 6.64, range: 67-96 years; 13 females, 5 males) participated in Experiment 1. All participants had binocular near visual acuity of 20/25 or better, with no differences in near or far acuity between groups (*p*s > .5). Two younger participants did not complete the contrast sensitivity test due to time constraints. The available data collected from 18 older and 16 younger adults showed that older adults had significantly lower contrast sensitivity at the highest spatial frequencies (12 and 18 cycles per degree or cpd) as measured using an Optec 6500P device (12 cpd: *t*(32) = 2.46, *p* = .019, *d* = 0.85; 18 cpd: *t*(32) = 3.22, p = .003, *d* = 1.11). Older adults scored higher on vocabulary than younger adults (*M* = 15.50, *SD* = 1.69; *M* = 12.90, *SD* = 2.18; *t*(34) = 3.93, *p* < .001, *d* = 1.31). We did not conduct a formal power analysis for Experiment 1, as its primary goal was to estimate the 75% point on the psychophysical function relating percent accuracy to luminance decrement in the target words. However, pilot data indicated a robust and steep relationship between pixel luminance decrement and accuracy, with a decrement of 11 pixels in the target letters yielding near-chance accuracy (∼50%) and a decrement of 44 pixels producing near-ceiling accuracy (∼100%).

#### Experimental Design

In a within-subject design, Experiment 1 manipulated the target word’s pixel luminance to identify the level for 75% accuracy for each age group. All trials used an unrelated prime. Participants completed 180 trials, with 45 trials at each of the four luminance decrements (11, 22, 33, and 44) chosen based on pilot data.

### Results

Figure 4 plots the mean probability of correctly identifying the target word as a function of the decrement in the mean pixel luminance of the target letters relative to the mean pixel luminance of the background noise for both younger (red) and older (blue) adults. The following psychometric function^1^ was fit to the data of both groups of participants.

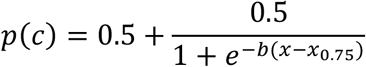

where *x* is the decrement in mean luminance of the target letters, 𝑥_0.75_ is the decrement in mean luminance corresponding^1^ to the probability of being correct = 0.75, and *b* determines the slope of the psychometric function. Figure 4 (left panel) shows that the psychometric functions fit to the two age groups fall very close to the points representing the obtained probabilities of being correct except for the decrement of 11-pixel values in the target word for older adults, which is lower than the guessing level of 0.5. The dashed red line (24.82 pixels) specifies the 75% accuracy target decrement for younger adults, and the solid blue line (26.01 pixels) specifies it for older adults. Hence, in Experiment 2, the target word’s pixel decrement was set to 25 for younger adults and 26 for older adults (for all three prime types).

**Figure 4.**
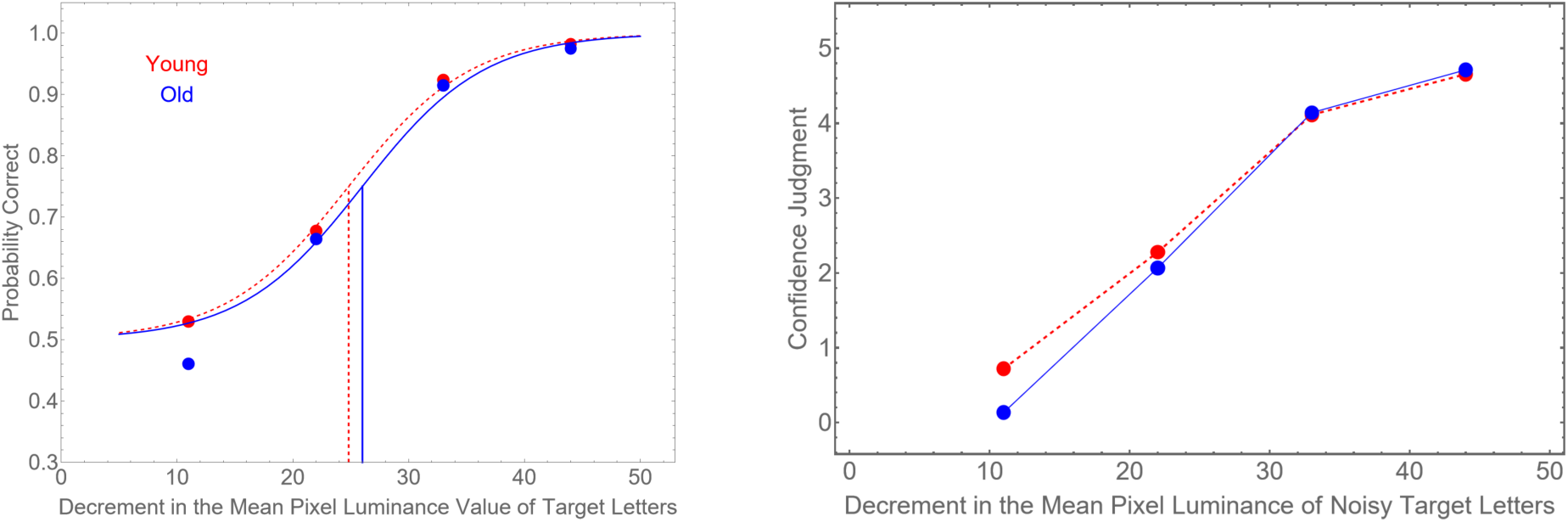
Accuracy and Confidence by Target Luminance. *Note*. (Left) Probability of correct identification in the unrelated prime condition as a function of target pixel decrement for younger (24.82, dashed red) and older (26.01, solid blue) adults. (Right) Mean confidence judgments for both groups as a function of pixel decrement (unrelated prime condition).

As shown in Figure 4 (right), confidence judgments were similar for most decrements (22, 33, and 44 pixels). However, the 11-pixel decrement showed a significant difference (p<.001, r_rb_ = -.82), with a large effect size (Mann-Whitney test, Cureton, 1956).

## Experiment 2

### Method

#### Participants

Thirteen younger adults (*M* = 21.20 years, *SD* = 2.08, 19-25 years; 12 females, 1 male) and fourteen older adults (*M* = 74.80 years, *SD* = 3.36, 68-79 years; 9 females, 5 males) initially participated in Experiment 2. One older adult did not complete the contrast sensitivity test due to time constraints. Compared to younger adults, older adults showed lower contrast sensitivity at higher spatial frequencies (12 cpd, *t*(24) = 3.01, *p* = .007, *d* = 1.2; 18 cpd, *t*(24) = 4.88, *p* < .001, *d* = 1.91) but showed no differences in binocular far or near visual acuities, or in performance accuracy at lower spatial frequencies (*p*s > .05). One younger adult did not complete the Mill Hill test due to a time constraint, older adults scored higher than younger adults (*M* = 15, *SD* = 2.54; *M* = 12.60, *SD* = 1.51; *t*(24) = 2.88, *p* = .008, *d* = 1.13). T-tests confirmed no significant demographic differences between Experiment 1 and Experiment 2 for both age groups (ps>.05).

One older adult was removed from the final analysis due to performance more than 6 SD below the mean, resulting in a final sample of 13 younger and 13 older adults for priming data analysis. A power analysis indicated that the power corresponding to the sample size used in this study (13 younger and 13 older adults) to detect an interaction effect comparable to the effect size reported in Experiment 1 from Jacoby et al., (2012; *η_p_^2^* = .24) was greater than 99% at *α* = .05. This power analysis was performed using G*Power 3.1 (Faul et al., 2007).

#### Experimental Design

Experiment 2 consisted of a single intermixed block with 60 unrelate prime trials, 60 Prime = Target trials, and 60 Prime = Foil trials. Unlike Experiment 1, the pixel decrements applied to the pixel luminance values of the target letters were fixed at 25 for younger adults and 26 for older adults.

## Results

### Accuracy of Target Word Identification as a Function of Prime Type

Younger and older adults’ unrelated prime performance is approximately the 0.75 target value (Figure 5, left). However, older adults outperform younger adults when the target equals the prime and underperform when the target equals the foil. When averaged across younger and older adults, average performance appears to be equivalent, independent of whether the prime is the same as the target or the same as the foil, and slightly higher than when the prime is unrelated to the foil (Figure 5, right).

**Figure 5.**
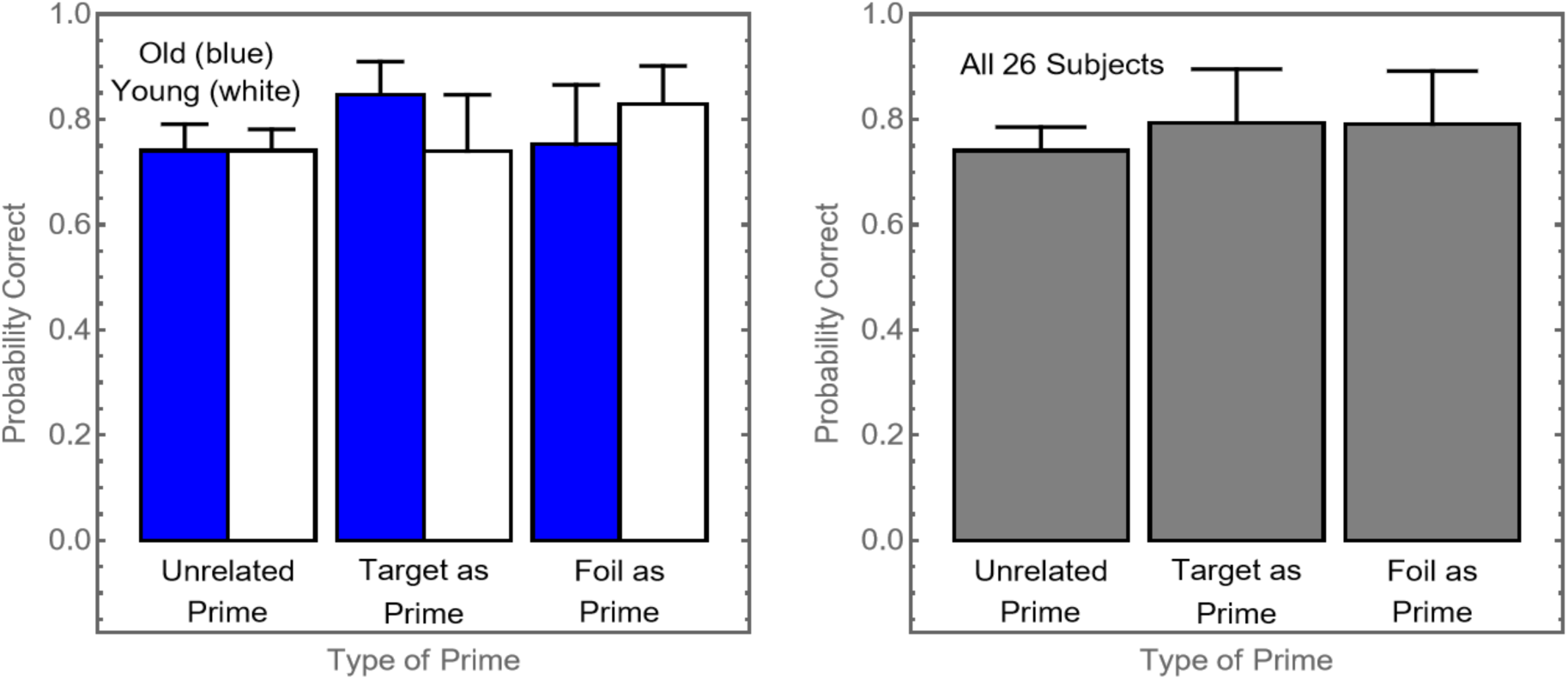
Accuracy by Prime Type and Age Group. *Note.* (Left) Probability of correct response as a function of prime type, plotted separately for older (blue) and younger (white) participants. (Right) Probability of correct response for all 26 participants across the three prime types. Error bars indicate 1 standard deviation. Standard deviation is used here, rather than standard error, to better visualize the differences in variance between conditions, as reported later in the Results.

A notable feature of these plots is that the SD of correct probability was smaller in the unrelated prime condition than in the two related conditions. Two-tail F-tests confirmed that the variance for all 26 participants was significantly lower in the unrelated condition compared to both the Prime = Target (*F*(25, 25) = .189, *p* < .005) and Prime = Foil (*F*(25, 25) = .193, *p* < .005) conditions. The variances of the two related conditions, however, did not significantly differ from each other (*F*(25, 25) = .98, *p* > .05). Due to the substantial variance difference between related and unrelated primes, non-parametric tests were used. A Mann-Whitney test comparing young and old adults collapsed across all three primes was not significant (*p* > .5). However, a 2-tailed Sign test showed a significant difference between unrelated and averaged related prime conditions (*p* < .001).

Two separate one-sample t-tests found no evidence that percent correct in Experiment 2 in the unrelated prime condition differed from 75% for either younger or older participants (*t*(12) = -0.82, *p* = .43, *d* = -0.23; *t*(12) = -.64, *p* = .53, *d* = -0.18). A paired-sample t-test found that no age difference (*M* = 74.1% for both groups, *SD* = 0.04 for younger adults, *SD* = 0.05 for older adults) in performance accuracy: *t*(24) = -0.01, *p* = .99. These results successfully validate that visual perceptual difficulty was statistically equated at the target 75% threshold.

A repeated-measures ANOVA examining the interaction between age group (2 levels: younger, older) and related priming conditions (2 levels: prime = target, prime = foil) on performance accuracy found a significant interaction (*F*(1, 24) = 10.49, *p* = .003, *η_p_^2^* = .30). Older adults outperformed younger adults in the prime = target condition (*M* = 0.85, *SD* = 0.06 vs. *M* = 0.74, *SD* = 0.11), while younger adults outperformed older adults in the prime = foil condition (*M* = 0.83, *SD* = 0.07 vs. *M* = 0.75, *SD* = 0.11). Neither the main effect of age group (*F*(1, 24) = 0.50, *p* = .48, *η_p_^2^* = .02) nor the main effect of priming (*F*(1, 24) = 0.008, *p* = .92, *η_p_^2^* = .0003) was significant.

### Signal Detection Analysis of Age-Related Differences in Response Bias

A signal-detection analysis, using the model described in Figure 1, was applied to the two related prime conditions (Prime = Target, Prime = Foil) to assess *d′* and *c*. In this analysis, the probability of a Hit corresponds to the Target = Prime condition, and the probability of a False Alarm (FA) corresponds to the Target = Foil condition. The model was fit to the individual data of all 26 participants. A two-tailed t-test comparing the mean *d′* values between younger (*M* = 1.67, *SD* = 0.42) and older adults (*M* = 1.79, *SD* = 0.37) did not reach significance (*t*(24) = -0.75, *p* = .46, *d* = -0.29). This result confirms the successful equating of perceptual sensitivity achieved via the individualized contrast settings. In contrast, a t-test of the response bias showed a significant age-related difference with a large and robust effect size (*t*(24) = 3.13, *p* < .005, *d* = 1.23). Younger adults showed a positive response bias (*M* = 0.15, *SD* = 0.25), whereas older adults showed a negative response bias (*M* = -0.17, *SD* = 0.28). This significant difference indicates that older adults require less perceptual evidence than younger adults to conclude that the target matches the prime word (Figure 6).

**Figure 6.**
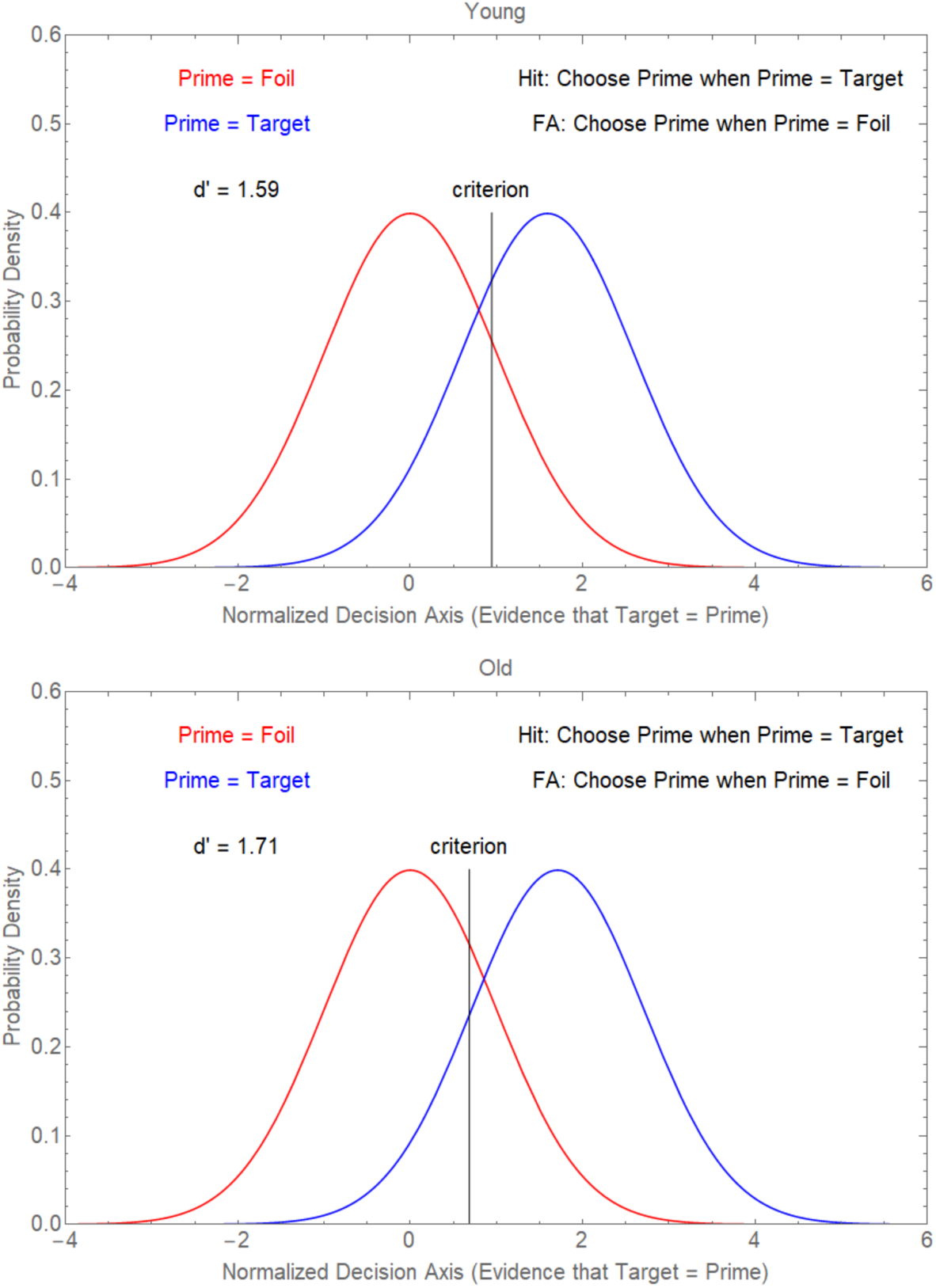
Age Differences in Response Bias (c) and Sensitivity (d′) *Note*. Similar to Figure 1, the x-axis denote that the Decision Axis is (evidence that Target = Prime). The blue distribution (Prime = Target) and red distribution (Prime = Foil) are separated by *d′* (Younger: 1.59; Older: 1.71). The area under the blue and red distributions to the right of the criterion specifies the probability of a Hit and a False Alarm (FA), respectively. The criterion (*c*) determines the decision boundary, with hits and false alarms measured by the area under the distributions to the right of c. While d′ is comparable across groups, the model fit shows younger adults have a positive *c* and older adults have a negative *c*. This demonstrates that older adults require less evidence than younger adults to conclude the target matches the prime.

Exploratory correlation analyses examined the relationships between response bias and demographic factors for each group. Neither age nor vocabulary score were correlated with response bias for younger adults (*r* = -.39, *p* = .18; *r* = .20, *p* = .52). Although neither correlation was found significant for older adults (*r* = -.41, *p* = .17; *r* = .53, *p* = .06), both correlation coefficients were moderate in strength.

### Confidence Judgments Across Priming Conditions and Age Groups

Figure 7 shows how the confidence ratings varied with Type of Prime and Age. A repeated measure 2 (age group: younger, older) by 3 (priming conditions: unrelated prime, prime = target, prime = foil) ANOVA revealed a main effect of condition (*F*(2, 48) = 27.90, *p* < .001, *η_p_^2^* = .54) and an interaction between age and condition (*F*(2, 48) = 4.08, *p* = .023, *η_p_^2^* = .15). However, there was no significant main effect of age (*F*(1, 24) = 0.61, *p* = .442, *η_p_^2^* = .03). Both age groups showed increased confidence from the baseline to the same and different priming conditions, with older adults demonstrating slightly greater increases in confidence (*M* = 2.59, 3.38, and 3.2 respectively) than younger adults (*M* = 2.56, 2.9, and 2.89 respectively).

### Modeling the Age by Prime-Type Interaction and the Role of Response Bias

The equivalence in the unrelated priming condition (Figure 5) and the crossover interaction (Target = Prime: Older > Younger; Target = Foil: Older < Younger) suggests an age-independent performance boost from related primes with an age-dependent bias. Averaged over groups, related primes boosted performance by 5.13% (*M_related_* = 79.23% vs. *M_unrelated_* = 74.10%). We quantified the contribution of response bias by calculating the average absolute Age × Prime-Type interaction effect (4.62%), suggesting response bias alone could account for differences in performance.

If response bias is indeed the major factor driving the performance differences in the related priming conditions, we could predict that the average performance of young subjects when the prime was equal to the target would be reduced by the average absolute size of the interaction effect (4.62%) from 79.23% to 74.62% and increased to 83.85% when the prime was equal to the foil. Meanwhile, the average performance of older subjects would be increased by 4.62% from 79.23% to 83.85% when the prime equals the target and decreased to 74.62% when prime equals the foil. The model’s predictions closely capture the observed interaction (Figure 8), strongly suggesting that differences in response bias drive the age effects in the lexical priming task.

## General Discussion

The current study was designed to investigate whether older adults’ increased reliance on contextual information is driven by sensory deficits or response bias. To separate these factors, we used a two-experiment design to first equate baseline perceptual performance across two age groups, then used a novel SDT analysis to examine the hypothesis that any remaining age-related differences in priming effect would be attributed to response bias. Our results largely supported this hypothesis, as Experiment 2 showed that with equated perceptual sensitivity, the age-related crossover in priming performance accuracy was attributable to a significant difference in response bias: older adults showed a negative bias, and younger adults showed a positive bias.

Hence, the priming effect of related words - whether the prime was a foil or a target - enhanced word identification accuracy in both younger and older adults. Priming occurs when prior exposure to a related stimulus facilitates the processing of a subsequent stimulus, often by activating shared or proximate representations in memory (Faulkner, 1983; Madden, 1988). For example, because a foil word shares semantic or phonological features with the target word, its presentation can activate both itself and the associated target in the observer’s mind, thereby facilitating recognition of the target word. Nevertheless, older adults exhibited a stronger reliance on consistent primes, aligning with conclusions from a meta-analysis by Laver and Burke (1993). This pattern also supports findings that older adults compensate for sensory and perceptual declines by increasingly relying on contextual information (Janse et al., 2014). Note that in real-world settings, contextual cues are typically consistent with the target (e.g., the gorilla example described in the introduction). Therefore, reliance on contextual cues, in general, is thus likely to be a quite adaptive strategy which allows older observers to optimize target identification under challenging viewing situations.

The SDT model in Figure 1 provides a clear mechanism for these results. In this 2ADFC task, a small shift in the response bias can produce the exact crossover interaction we recorded in Experiment 2. Our model described above demonstrates that a slight negative shift in response bias boosts hit rates in the prime = target condition, while increasing errors in the prime = foil condition. Conversely, the slight positive shift in response bias produces the opposite pattern. This alignment strongly suggests that the observed age-related differences in priming, including the “false seeing” phenomenon reported in previous studies, is a direct consequence of older adults having a negative response bias, whereas younger adults have a positive response bias.

Consequently, our results attribute these age differences to response bias: younger adult’s positive bias suggested that they required more evidence to reach a prime-match conclusion, whereas older adults’ negative bias made them more accepting of a match, therefore, becoming more susceptible to foil errors. Notably, the magnitude of this difference was large (*d* = 1.23; Cohen, 1988). This response bias is most likely a result of older adults’ greater reliance on context, relative to younger adults, when sensory factors render perception difficult. Jacoby et al., (2012) refers to this effect as “false seeing” suggesting that the output of perceptual processing in older adults in such situations is perception of the prime as the same as the target. In contrast, the current work identifies a response bias mechanism responsible for this phenomenon, rather than actual perception of the wrong target.

In both models of the results (response-bias versus false-seeing) we would expect, given that the stimuli were equated for perceptual difficulty, that older and younger adults would be equally confident concerning their identification responses in the presence of an unrelated prime, but that older adults would be more confident than younger adults when the prime was related to the target (Figure 7). However, if the ‘false-seeing’ interpretation is correct, we would expect participants’ tendency to identify the target as the prime to remain relatively unaffected by common manipulations known to influence response bias, such as feedback, stimulus frequency, or motivational incentives. Future research should directly examine whether older adults’ ‘false seeing’ tendencies are modifiable by any factors that are known to affect response bias.

**Figure 7.**
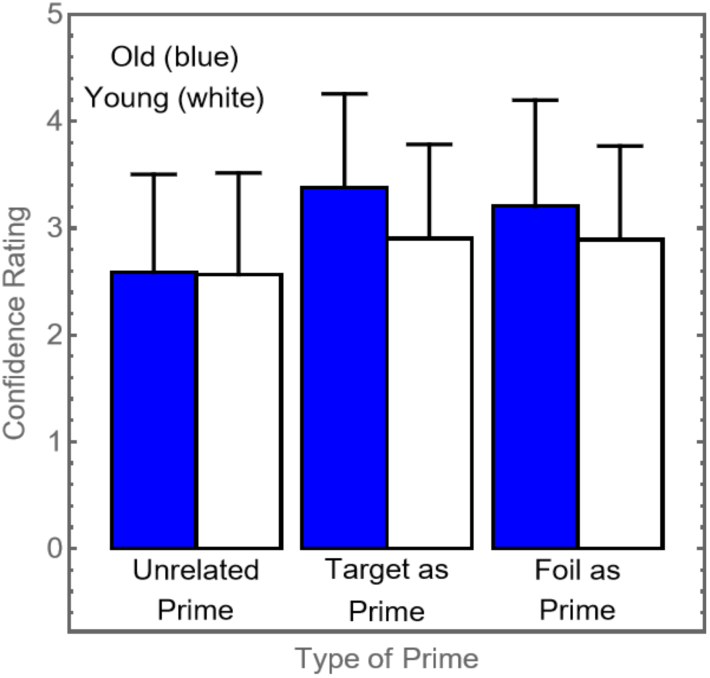
Confidence by Prime Type and Age Group. *Note.* Confidence as a function of the type of prime, plotted separately for older (blue) and younger (white) participants. The error bars indicate 1 standard deviation above the mean.

**Figure 8.**
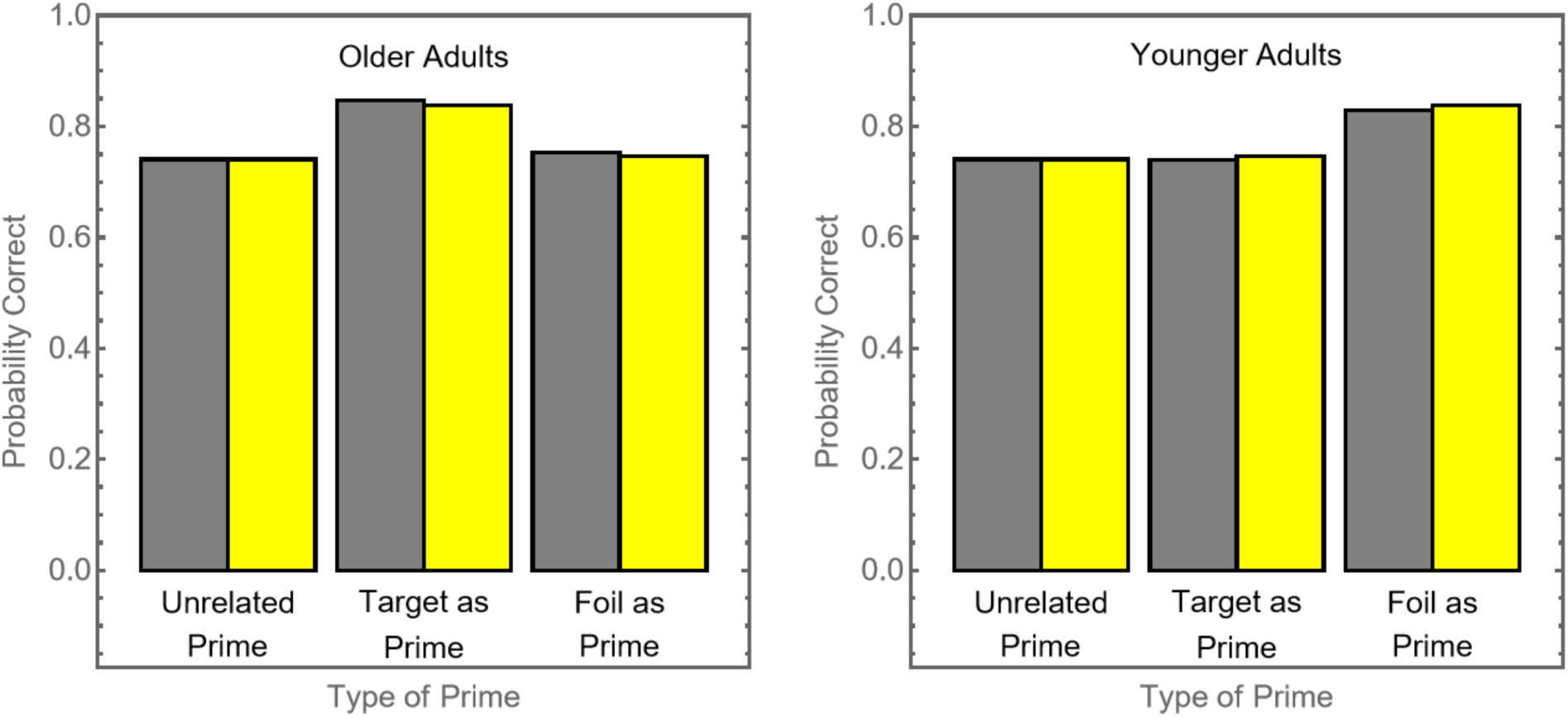
Model Fit of Priming Effects by Age Group. *Note*. Model Fit of Age × Priming Interaction. Probability of correct word identification for older (left) and younger (right) adults across three priming conditions. The model’s predictions (yellow) closely align with the observed data (grey), demonstrating that a change in response bias captures the Age × Priming interaction.

The analysis showed a large effect size for age differences in response bias, indicating that a shift in decision-making strategy, rather than solely perceptual decline, substantially drives the observed age differences in the priming effect. Future studies examining age and visual deficits should consider integrate Signal Detection Theory analysis to quantify and account for the contribution of response bias on behavioural performance.

A handful of studies in the literature find that older adults exhibit a more conservative response bias in simple and complex visual perception tasks but that this pattern can shift depending on task demands and paradigm (Thapar et al., 2003). In Jacoby et al. (2012), older adults outperformed younger adults by 28% when the prime was the same as the target but underperformed them by 17% when the prime was the same as the foil. In our current study, where we carefully controlled for age-related differences in performance when the prime was unrelated to the target, older adults outperformed younger adults by 11% when the prime was equal to the target and underperformed them by 8% when the prime was equal to the foil. These differences in “false seeing” between our and the Jacoby et al. study could be due to the lability of response biases, meaning it could be influenced by various experimental design factors, such as feedback, task instructions, and contextual cues. The fewer constraints placed on the experimental design; the more variable response bias may become. Hence, differences in experimental constraints and design could contribute to differences in the observed false seeing effect. For example, providing feedback as to the correctness of a response, using an open-ended recall task instead of limiting responses to two choices, or providing monetary rewards for correct identifications or costs for incorrect identifications, could change response biases and, consequently, affect the size of the false seeing effect. Hence, age-related differences in the strategic adjustment of response biases, initiated by testing conditions, feedback, and outcome consequences, rather than fundamental differences in the early stages of perceptual processing, could be the basis of ‘false seeing’ in older adults.

Despite low power, our analysis revealed a noteworthy correlation within the older adult group. Older adults with a larger vocabulary size showed a younger-like response bias. Future research should consider a wider battery of cognitive measures (e.g., executive function, working memory capacity) and linguistic measures (e.g., verbal fluency test, a naming task) to determine if this relationship is specific to verbal knowledge or reflects generic cognitive reserve in older adults. Including middle-aged participant could also help clarify the development trajectory of this relationship between cognitive abilities and decision bias.

## Limitations

Although our findings provide clear insights into age-related differences in contextual reliance, several limitations must be acknowledged. First, the sample sizes for both experiments were relatively small, and the sample size for the first experiment was determined by our previous study and a pilot study without formal power analysis. However, psychometric functions relating stimulus contrast to recognition performance typically show very steep slopes, meaning small contrast changes lead to large variations in accuracy. Thus, even moderate sample sizes can yield highly precise estimates of contrast thresholds corresponding to approximately 75% accuracy. Nevertheless, future research using larger samples could further validate and refine these estimates from SDT. Second, the cross-sectional nature of our design and the use of extreme age groups (younger vs. older adults) could be subject to limitations regarding potential cohort effects and generalizability across the adult lifespan (Freund & Isaacowitz, 2014).

Initially, we planned to recruit middle-aged adults to provide a continuous developmental perspective; however, this was not possible due to the onset of the COVID epidemic in 2020 which forced the closing of the lab, and subsequent retirement of the principal investigator in 2022 before the end of the epidemic, which made the closure permanent. Furthermore. Our sample was not balanced for gender: for example, including only 1 younger male participant in Experiment 2 could limit the generalization of our findings, which is especially important given reported sex differences in lexical priming effects (Bermeitinger et al., 2008). Finally, we did not examine the role of cognitive and linguistic covariates, such as working memory capacity, on contextual reliance and response bias. Future research incorporating these variables could provide a more fine-grained picture of individual differences in the response strategies employed by older adults when processing ambiguous perceptual information.

## Conclusion

This study examined age-related differences in contextual reliance during visual word recognition by equating perceptual difficulty and applying SDT. We found that the increased reliance on contextual cues in older adults was primarily driven by response biases rather than perceptual sensitivity differences, highlighting differences in the cognitive decision-making strategies used by younger and older adults. These results emphasize the importance of distinguishing between perceptual and cognitive contributions to age-related changes in visual cognition, providing insights for interventions aimed at supporting older adults in perceptually challenging environments.

## Footnotes

1 An iterative process was used to find the minimum value of Pearson’s Chi Square between the predicted and obtained number of correct and incorrect responses at each of the 4 levels of *x*. Because there were 45 trials at each of the 4 values of *x* for each of the 18 adults in a group, the predicted number of correct trials at a given value of *x* is equal to 45*18*p(c).

## Notes

### Competing Interest Statement

The authors have declared no competing interest.

